# A novel, ssDNA Bidnavirus in the giant freshwater prawn *Macrobrachium rosenbergii*

**DOI:** 10.1101/2022.09.25.509311

**Authors:** Warachin Gangnonngiw, Malinee Bunnontae, Pattanapon Kayansamruaj, Saengchan Senapin, Jiraporn Srisala, Timothy W. Flegel, Kanokpan Wongprasert

**Author notes:** **Corresponding authors** K. Wongprasert.

## Abstract

A purported parvovirus producing circular, eosinophilic inclusions in the nuclei of tubule epithelial cells of the hepatopancreas (HP) of the giant freshwater prawn *Macrobrachium rosenbergii* (*Mr*) was first reported from Malaysia in 1994. In 2009, similar inclusions in *Mr* from Thailand were confirmed to contain single-stranded DNA (ssDNA). In 2021, *Mr* samples showing similar HP inclusions were reported to give positive PCR test results for decapod *Hepanhamaparvovirus* (DHPV) using universal DHPV primers designed from many viral isolates of 3 penaeid shrimp species. However, the matching DNA probe revealed positive *in situ* hybridization (ISH) results with host HP nuclei only, and not with the intranuclear inclusions. At the time, the negative ISH result with the intranuclear inclusions has not been further elucidated. Here we describe metagenomic analysis and *de novo* assembly from 2016 samples of *Mr* with similar intranuclear inclusions that showed negative PCR test results using the DHPV universal primers. Genomes identified through metagenomics have revealed a novel ssDNA genome sequence of 6,723 bases that was confirmed by a single PCR amplicon. ISH using specific probes derived from this genome sequence demonstrated the ssDNA presented in both grossly normal nuclei and in the intranuclear inclusions. Tests with archived DNA from the samples used in the 2021 publication yielded positive PCR results for both DHPV and the novel sequence, suggesting that the shrimp had dual infections. Phylogenetic analysis of the novel sequence revealed no relationship to any known DHPV type. Instead, the sequence most closely matched the polymerase B gene in the V1 fragment from *Bombyx mori bidensovirus* 2 (BmBDV2) is characterized by 2 ssDNA genome fragments and has been classified in the family *Bidnaviridae*. The 2 BmBDV2 fragments V1 and V2 each produce their own independent capsid protein genes and virions. In contrast, the new *Mr* virus is independent with a single 6,723 base ssDNA genome, and we propose the name Macrobrachium hepatopancreatic bidnavirus (MHBV).

## 1. INTRODUCTION

In 1994 a putative parvovirus was described in the hepatopancreas of post-larvae (PL) of the giant freshwater prawn *M. rosenbergii* (*Mr*) in Malaysia (Anderson et al., 1990) based on light and electron microscope observations of circular, eosinophilic to magenta, intranuclear inclusions in tubule epithelial cells of the hepatopancreas (HP). Similar inclusions were reported from PL of *Mr* in Thailand in 2009, and they were shown to contain single-stranded DNA (ssDNA) (Gangnonngiw et al., 2009), supporting the proposal that they might be caused by a parvovirus. However, *in situ* hybridization (ISH) tests on similar tissue from Malaysia using a DNA probe developed from a parvovirus in the HP of the penaeid shrimp *Penaeus chinensis* (Lightner et al., 1994; Mari et al., 1995) gave negative test results. Subsequent tests using *Mr* samples from Thailand with both PCR and ISH methods designed for a hepatopancreatic parvovirus in the penaeid shrimp *Penaeus monodon* (Phromjai et al., 2002) also gave negative results. Due to sample limitation, a genetic study could not be carried out. In 2016, more *Mr* samples with similar intranuclear inclusions were obtained and DNA extracted from these samples was sent for whole genome shotgun sequencing (WGS). This report describes the results of sequencing and analysis plus testing by PCR and ISH.

In parallel with work described herein, a different laboratory developed a potential universal PCR detection method for the group of closely related hepatopancreatic parvoviruses that infect several penaeid shrimp (*P. chinensis* and *P. monodon* in the previous paragraph). These viruses produce ovoid, magenta to basophilic, intranuclear inclusions in the HP (Srisala et al., 2021). These penaeid shrimp viruses were originally called hepatopancreatic parvoviruses (HPV), the name was later changed to the genus *Hepanvirus*, in the sub-family *Densovirinae* (DHPV) in the sub-family *Hamaparvovirinae* (Pénzes et al., 2020). Recently, Srisala et al. (2021) used their universal PCR detection method with an independent set of *Mr* samples and obtained a positive result for DHPV with a PCR amplicon of 100% sequence identity to the control amplicon, indicating the presence of a type of DHPV in the *Mr* samples. However, the ISH test results revealed that the HP cell nuclei of normal morphology gave positive ISH results for DHPV but that the intranuclear inclusions in the same samples did not. The authors reported that they could not explain the negative ISH result with the intranuclear inclusions, but that the positive results in the superficially normal HP cell nuclei supported the proposal that the *Mr* samples were infected with a new type of DHPV. Work on sequencing the whole genome for confirmation of this DHPV type is underway. However, the negative ISH results raised the possibility that the intranuclear inclusions arose from something other than a type of DHPV, and possibly a novel virus.

Here, we describe the discovery of a novel virus in *Mr* that is distinct from DHPV and is linked by positive ISH results to the mystery intranuclear inclusions described above. We call it Macrobrachium hepatopancreatic bidnavirus (MHBV) and we show that it sometimes occurs together with DHPV in dual infections. Hatchery mortality of PL associated with MHBV intranuclear inclusions is of relatively rare occurrence, so there are few publications related to it.

## 2. MATERIALS AND METHODS

### 2.1. Source of samples

Our interest in this work began in 2009 (Gangnonngiw et al) upon the receipt of an *Mr* hatchery incident involving mortality of post larvae (PL). Histological examination revealed the presence of magenta, intranuclear inclusions in HP tubule epithelial cells of some of the specimens. Only a few archived paraffin blocks remained for possible further work by histological analysis and *in situ* hybridization. However, the primary samples used herein consisted of grossly normal *Mr* post larvae (PL) obtained from a different hatchery in Thailand in October 2016. The HP were removed from some PL and divided into two parts. One part was fixed in 4. 5% glutaraldehyde for transmission electron microscopy, and the other was kept at - 80 ºC for DNA extraction. The remainder of the samples were fixed for histological analysis and ISH analysis.

A second set of samples consisting of the archived paraffin blocks from 2009 (Gangnonngiw et al., 2009) were also used for histological examination including ISH assays. A third set of *Mr* samples of importance to this study consisted of PL collected in 2017 and 2018 from the same hatchery as our primary samples but used for the development of a universal PCR method for DHPV published by Srisala et al. (2021). From that study, only archived DNA extracts remained for further analysis. However, photomicrographs in the 2021 publication clearly revealed that the HP intranuclear inclusions gave negative ISH test results with the DHPV-specific probe developed. Using 10 archived DNA samples from the 2021 study, PCR analyses for both DHPV and MHBV were carried out, but no material remained for histological analysis and MHBV-ISH assays.

Finally, in 2022 a fourth set of 43 normal PL samples was obtained from the same source hatchery as the primary samples described herein 2016. These were tested for both DHPV and MHBV by PCR. Due to the limited number of samples and the discovery time course, all tests could not be carried out with all sample sets. To avoid confusion due to the disjointed variety of samples and sampling times, **Table 1** shows a summary of the sample sets together with tests to which they were subjected and their results. Handling of animals and the study protocol was approved by the Mahidol University Animal Ethics Committee (Protocol No. MUSC61-055-457).

**Table 1.** Summary of the *Mr* PL sample sets included in this study and results of various tests carried out with each set. +, positive result; -, negative result; n/a, not applicable; ND, not done; * no dual positive samples; ** ISH probe for DHPV type in *Penaeus chinensis*

| Test or observation | Sample sets and years sourced |  |  |  |  |
| --- | --- | --- | --- | --- | --- |
|  | Previous publication 1994 | Set one 2009 | Set two 2016-2018 | Set three 2016 | Set four 2022* |
| Associated PL mortality | n/a | + | + | - | - |
| Intranuclear inclusions by histology | + | + | + | + | - |
| ssDNase digestion | ND | + | ND | ND | ND |
| ISH for Pchin-DHPV** | - | ND | ND | ND | ND |
| PCR for Pm-DHPV | ND | - | - | - | ND |
| ISH for Pm- DHPV | ND | - | - | - | ND |
| Universal- DHPV PCR | ND | ND | + | - | +2/43* |
| ISH for U-DHPV in nuclei | ND | ND | + | - | ND |
| ISH for U-DHPV in intranuclear inclusions | ND | ND | - | - | ND |
| PCR for MHBV | ND | ND | + | + | +1/43* |
| ISH for MHBV- in nuclei | ND | ND | ND | + | ND |
| ISH for MHBV in intranuclear inclusions | ND | ND | ND | + | ND |
| Reference | Lightner et al. 1994 | Gangnonngiw et al. 2009 & this publication | Srisala et al., 2021 & this publication | This publication | This publication |

### 2.2 Histological examination

Giant freshwater prawn post larvae of *M. rosenbergii* were fixed with Davidson’ s fixative and processed for routine histology using hematoxylin and eosin (H&E) staining as described by Bell & Lightner (1988). These paraffin-embedded tissues were also used to prepare sections for ISH assays.

### 2.3 Transmission electron microscopy (TEM)

Hepatopancreatic tissues of approximately 1 mm^3^ were fixed for 2 h at 4°C with 4. 5% glutaraldehyde in 0.1 M phosphate buffer, pH 7.4. The fixing solution was removed, and samples were postfixed in 1% (w/v) osmium tetroxide in 0.1 M phosphate buffer at pH 7.4 for 2 h, washed twice with distilled water, dehydrated in a graded series of ethanol solutions from 50 to 100%, followed by 2 immersions in 100% propylene oxide. Tissues were embedded in Epon-812 resin by successive 1 h infiltrations of 1:1 and 2:1 resin:propylene oxide followed by 100% resin. The tissue blocks were then polymerized by incubating at 70°C for 48 h in fresh 100% epoxy resin. To pre-screen samples for TEM, semi-thin sections were stained in toluidine blue. Then, blocks of samples with intranuclear inclusions were used to prepare thin sections stained in 2% (w/v) uranyl acetate and 0.3% (w/v) lead citrate solutions. The sections were viewed and photographed under a Hitachi H7100 electron microscope at 100 kV equipped with a digital camera.

### 2.4 DNA extraction

DNA was extracted using a QIAamp DNA mini kit (QIAGEN), and 10 µl of 25 mg/ml RNaseA was added during cell lysis. DNA degradation degree and potential contamination were monitored on 1% agarose gels and purity (OD260/ OD280, OD260/ OD230) was checked using a Nano Photometer^®^ spectrophotometer (IMPLEN, CA, USA). DNA concentration was measured using Qubit^®^ dsDNA Assay Kit in Qubit^®^2.0 Fluorometer (Life Technologies, CA, USA). OD values were between 1. 8∼2. 0, and DNA contents above 1 µg were used to construct a library for metagenomic sequencing.

### 2.5 Whole-genome shotgun metagenomic sequencing (WGS)

DNA was submitted to Getz Healthcare (Thailand) Ltd. for Qubit™ fluorometric analysis, library preparation with Nextera XT kit and high throughput DNA sequencing using the Illumina HiSeq 150× paired-end system. Low-quality sequences were trimmed using Trimmomatic v0.39 (Bolger, Lohse, and Usadel 2014).

### 2.6 Bioinformatics analysis

#### 2.6.1 De novo metagenome assembly and isolation of the viral genome

Trimmed reads were assembled using MEGAHIT v1. 2. 8 with the default setting (Li et al., 2015). Contigs (with a length of ≥ 500 bp) were then submitted to Kaiju web-based for taxonomy prediction of each contig by searching against the *nr* database for Bacteria, Archaea and Viruses (Menzel, Ng, and Krogh, 2016). Contigs with known identity as flagged automatically by Kaiju were subsequently subtracted from assembly to obtain contigs of unknown identity. These contigs were used to generate a local blast database in Blast2GO program and then blast (tblastn) using CDSs of *Bombyx mori* bidensovirus (BmBDV) (accession no. NC_020928) as query sequences (Götz et al., 2008). Contigs hit by both viral genomes were used for further investigation.

#### 2.6.2 Virus genome analysis and phylogenetic analysis

The genome retrieved from the previous step was annotated using the virus annotation mode of Prokka v1.14.0 (Seemann 2014). The identity and conserved domain of the viral ORFs were then identified using blastx against nr and Swiss-Prot databases, respectively. Inverted repeats present in the genome were predicted using EMBOSS einverted web tool with the minimum threshold of 20 and non-mismatch setting (Rice, Longden, and Bleasby 2000).

As it turned out, there was only one significant nucleotide translated protein (blastx) hit for any reading frame for the whole genome assembly of 6,723 bases. This was characterized by the presence of a highly conserved DNA polymerase B (PolB) domain. Thus, the only rational phylogenetic analysis was focused solely on this conserved sequence using Clustal Omega software.

### 2.7 PCR method

Preliminary tests were carried out with all DNA extracts using the proposed universal primers (DHPV-U primer) for detection of DHPV as previously reported from *Mr* in Thailand (Srisala et al., 2021). Detection of DHPV was also performed by using two other methods. One was a 1-step PCR method that employed primers HPV441F & R (product size 441 bp) (Phromjai et al., 2002) and the other was a semi-nested PCR method modified from Umesha et al. (2006) using primers HPV-VP-F & R and HPV-FN & HPV-VP-R (product sizes 449 bp and 214 bp respectively). In addition, a new PCR method with an amplicon of 392 bp was designed using the Primer3 (v.0.4.0) program based on the sequence of the new virus found in *Mr* by metagenomic analysis in this study. Amplification was performed with Taq DNA polymerase (GeneDireX^®^), primers left (5′-GCA TTA ATG GAT TGG GAA GG-3′), and primer right (5′-TCG ATG TCT GGA TGA CCG TA-3′). Temperature conditions comprised denaturation at 94 °C for 5 min followed by 35 cycles of 94 °C for 30 s, 53 °C for 30 s, and 72 °C for 2 min, followed by a final extension step at 72 °C for 5 min. Size examination was performed by electrophoresis on 1. 5% agarose gel. The beta-actin gene was used for internal control of the PCR method (Lu et al., 2006).

To amplify the whole genome product of 6,723 bases in one RT-PCR reaction, KOD FX Neo Taq DNA polymerase (Toyobo, Japan) was used with only one strand of primer derived from the forward inverted repeat sequence (5′-TTA TCC CCC TTT TCC ACA AAT AA -3′) complementary to that at the end of the genome. The temperature of denaturation was at 94°C for 2 min followed by 35 cycles of 98°C for 10s, 50°C for 30s, and 68°C for 3. 50 min, followed by a final extension step at 68°C for 7 min. Size examination was performed by electrophoresis on 0.8% agarose gel.

### 2.8 *In situ* hybridization (ISH)

A labeled DNA probe for the new virus derived from *Mr* was prepared using a commercial PCR DIG-labeling mix (Roche)according to the manufacturer’s instructions. Primers used for labeling the probe were the same as those used for the PCR method above (product size 392 bp). The negative control or negative probe for this experiment was labeled amplified DNA from the fish virus (TiLV). This non-target, negative-control DNA probe was used instead of a no-probe control to reveal any non-specific DNA binding such as sometimes occurs, for example, with the crustacean cuticle. Paraffin-embedded tissues sectioning and prehybridization protocols were performed according to Srisala et al. (2021). DIG-labeled probe (150 ng) was added to each sectioned slide. Negative controls consisted of adjacent tissue sections probed with the fish virus (TiLV) probe. After probe addition at 42°C overnight, the slides were washed, and the subsequent procedure was carried out as previously described (Srisala et al., 2021).

## 3. RESULTS AND DISCUSSION

### 3.1. Analysis of our primary 2016 specimens (sample set 3)

Histological investigation of hepatopancreatic tissue confirmed typical signs of eosinophilic to magenta, intranuclear inclusions characteristic of the target virus in the *Mr* samples. Previous reports and histopathology suggested that these inclusions in *M. rosenbergii* might arise from what are now called hepanhamaparviruses in penaeid shrimp (Flegel 2006). All 12 of the fixed *M. rosenbergii* specimens showed these intranuclear inclusions with H&E staining and with toluidine blue staining in semi-thin sections (**Fig. 1)**. Based solely on microscopic morphology and H&E staining, the intranuclear inclusions appeared to closely match those previously reported in *M. rosenbergii* from Malaysia (Anderson et al., 1990, Lightner et al., 1994) and from Thailand (Gangnonngiw et al., 2009; Srisala et al., 2021).

**Figure 1.**
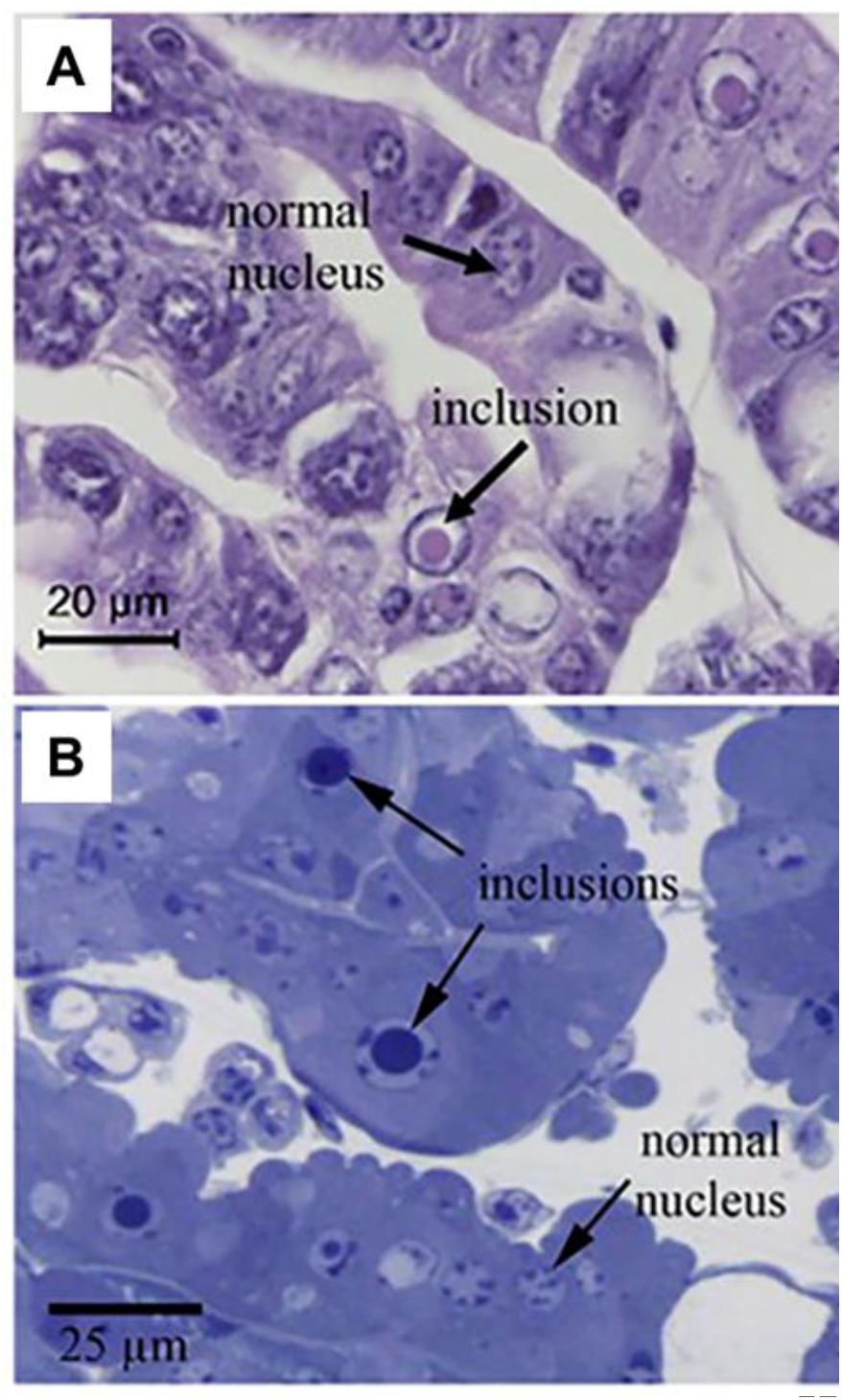
Example photomicrographs of *Mr* intranuclear inclusions by H&E and toluidine blue staining. (A) H&E-stained tissue section showing circular, magenta, intranuclear inclusions in HP tubule epithelial cells. (B) Semi-thin, toluidine blue-stained tissue section showing the same type of inclusions

Using the same block as for the semi-thin section shown in **Fig. 1**, thin sections were prepared for transmission electron microscopy (TEM). Virions could be seen in the intranuclear inclusion area (**Fig. 2**). They resembled those previously reported from *Mr* in Thailand (Gangnonngiw et al., 2009) but differed somewhat from those previously reported from Malaysia (Anderson et al., 1994) which showed some virions encased with a thick electron-dense structure.

**Figure 2.**
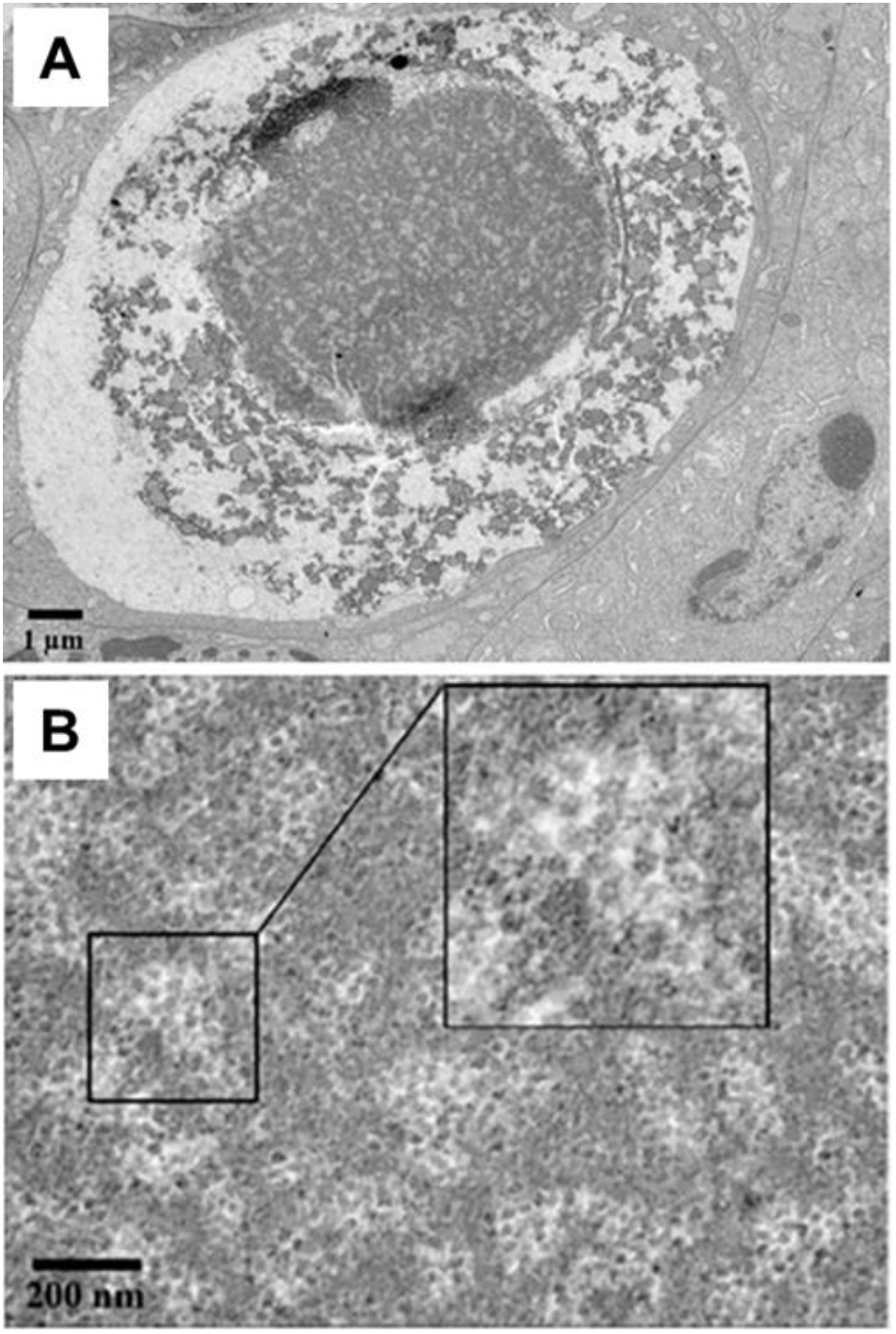
Transmission electron micrographs of an intranuclear inclusion in a hepatopancreatic tubule epithelial cell from an *Mr* post larva. (A) Low magnification electron micrograph. (B) High magnification micrograph with a magnified insert (Hitachi, HT7700).

For the aquaculture industry, those working on the development of specific pathogen-free stocks of *Mr* previously had to rely on histological investigation to test and monitor specimens for these viral inclusions due to the absence of genome sequence information that would allow for PCR testing. In addition, there was some doubt regarding the identity of the virus in these inclusions due to negative PCR test results and lack of ISH signals using methods developed for detection of DHPV (Srisala et al., 2021).

### 3.2 Negative PCR results obtained using DHPV primers

1. DNA samples from the HP of *Mr* postlarvae were used as templates for PCR tests performed using two methods. One was a 1-step PCR method that employed primers HPV441F & R (product size 441 bp) (Phromjai et al., 2002) and the other was a semi-nested PCR method modified from Umesha et al., 2006 using primers HPV-VP-F & R and HPV-FN & HPV-VP-R (product sizes 449 bp and 214 bp respectively). Both were designed to detect the DHPV previously reported from *Penaeus monodon* in Thailand. The results for both methods were negative (not shown) and indicated that the 2016 *Mr* samples (Set 3) were not infected with the same DHPV previously reported from *P. monodon* in Thailand. Subsequent tests with the same DNA samples using a recently published universal PCR method for DHPV (Srisala et al., 2021) also gave negative PCR test results with sample Set 3 (Fig. 8, lane 11). Due to the lack of positive test results using DHPV primers, it was necessary to carry out a metagenomic analysis to try to obtain genomic information from the Thai samples that showed DHPV-like intranuclear inclusions but gave negative test results using DHPV PCR detection methods.

### 3.3 A novel virus revealed in *Mr* by metagenomic analysis

WGS analysis was able to generate 10 Gb of raw reads data. Since full genomic data for *Mr* was not available in the public database, all obtained raw reads, including host-associated reads, were necessarily subjected to downstream bioinformatics analyses. Based on MEGAHIT assembler, a total of 458,696 contigs (>500-bp-long) were generated. Among these, 98% were described as unclassified origin by the Kaiju web program and only 0. 04% were characterized as virus-like sequences (0. 02%, 0. 01%, and 0. 005% were dsDNA, retro-transcribing, and ssRNA viruses, respectively), whereas no ssDNA viruses were detected. On the other hand, blasting against ‘unclassified’ contigs using densovirus CDSs as query sequences showed that non-structural and structural proteins gave hits for contigs with e-values of 2E-08 and 5E-05. The average coverage depth for one contig of 6,723-bp with 30.22 GC% was 299.7×. According to blast analysis (data not shown), a portion of ORF4 contained a DNA polymerase domain that gave high similarity to those reported from several members of the family *Bidnaviridae*, including Tasmanian devil feces bidnavirus, Bombyx mori bidnavirus (BmBV) and bidnavirus-like culex mosquito virus. Similar to other viruses of the family *Bidnaviridae*, the sequence harbored inverted repeat sequences called terminal complementary repeats (TCR) at each terminus of the 6,723-bp contig. PCR was used to confirm the metagenomic assembly sequence using primers designed based on the metagenomic data. A positive amplicon band of 6,723 bp was obtained by PCR (**Fig. 3A**) and sequencing confirmed that it matched the contigs produced from the metagenomic data.

**Figure 3.**
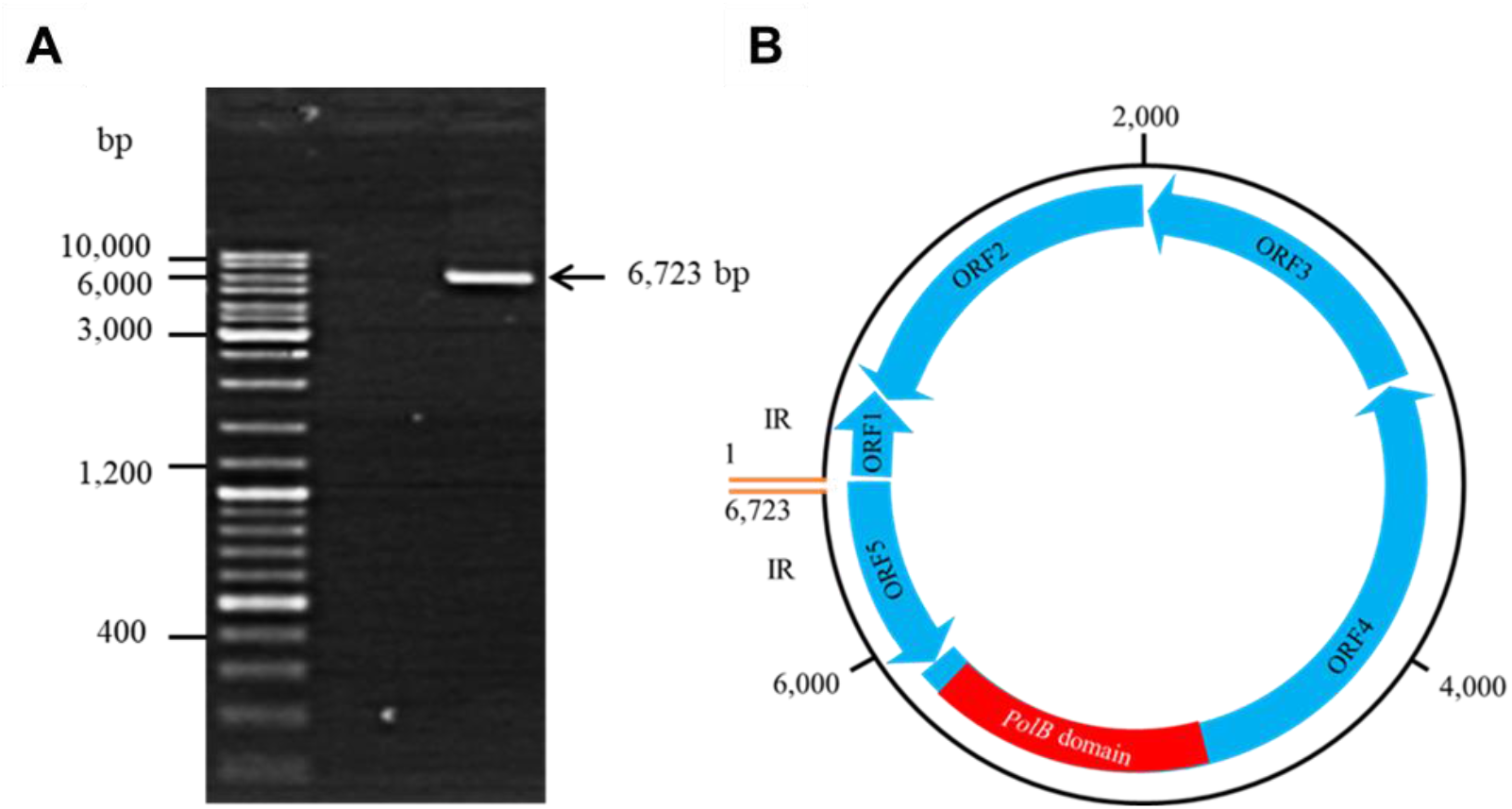
MHBV genome. (A) Whole DNA strand amplification of *Mr* hepatopancreas from post larvae (B) Tentative genome map based on the metagenomic assembly and using the BmBDV-V1 as the reference sequence. The ORF are indicated by light blue boxes, and the arrows represent the direction of transcription. Terminal inverted repeats (IR) are indicated as orange lines. The scale shows nucleotide positions corresponding to the genome map.

A schematic representation of the ‘panhandle’ structure (Hu et al., 2013) of the putative viral genome sequence based on reference to BmBDV-V1 is shown in **Fig. 3B**. A description of its putative ORFs is given in **Table 2**.

**Table 2.**
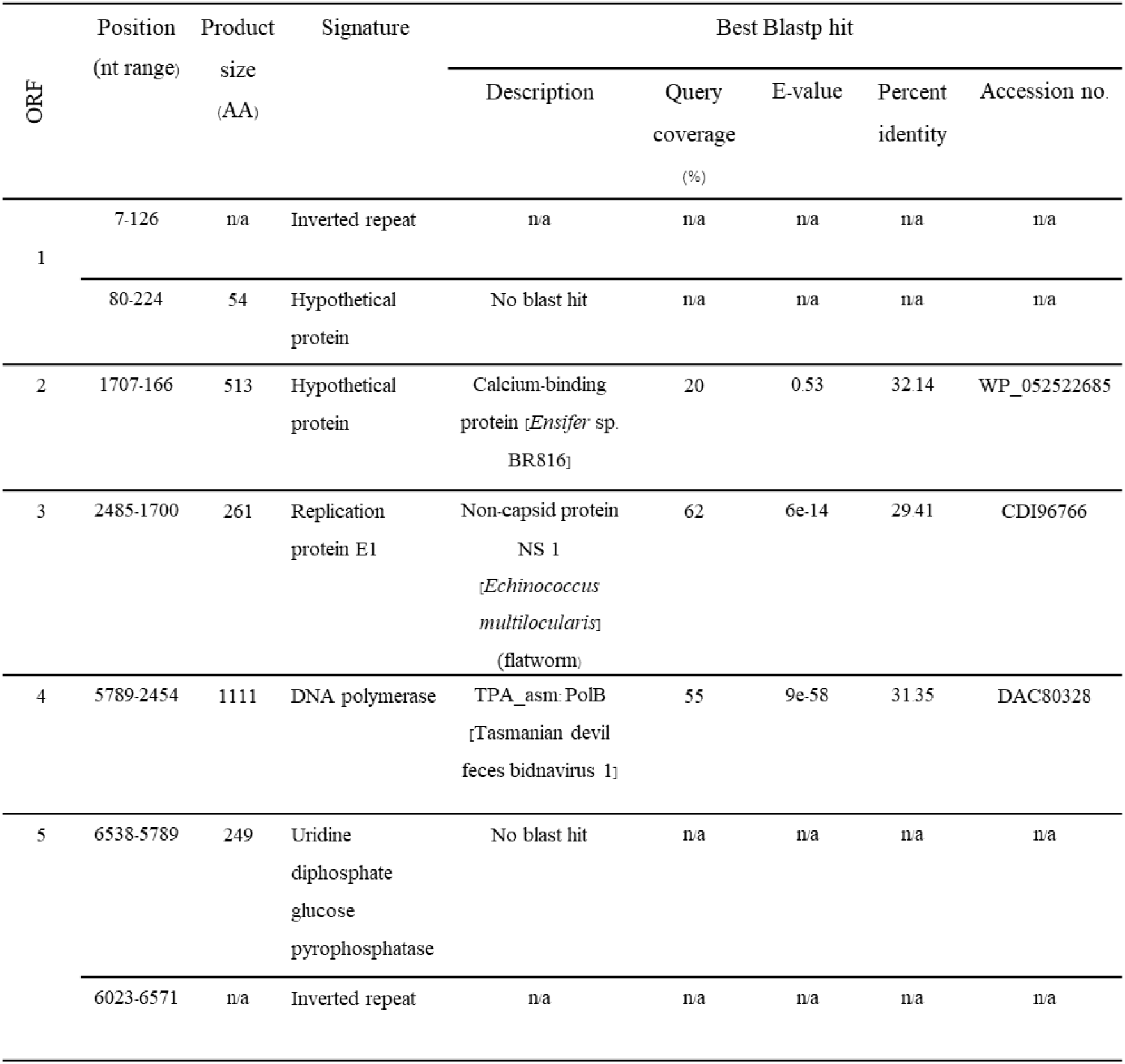
Genome annotation of the current MHBV sequence. Predicted function was determined based on Blastp search against Swiss-Prot database with an expected threshold of 10. n/a, not applicable.

| ORF | Position<br>(nt range) | Product<br>size<br>(AA) | Signature | Best Blastp hit |  |  |  |  |
| --- | --- | --- | --- | --- | --- | --- | --- | --- |
|  |  |  |  | Description | Query<br>coverage<br>(%) | E-value | Percent<br>identity | Accession no. |
| 1 | 7-126 | n/a | Inverted repeat | n/a | n/a | n/a | n/a | n/a |
|  | 80-224 | 54 | Hypothetical<br>protein | No blast hit | n/a | n/a | n/a | n/a |
| 2 | 1707-166 | 513 | Hypothetical<br>protein | Calcium-binding<br>protein [*Ensifer* sp.<br>BR816] | 20 | 0.53 | 32.14 | WP_052522685 |
| 3 | 2485-1700 | 261 | Replication<br>protein E1 | Non-capsid protein<br>NS 1<br>[*Echinococcus*<br>*multilocularis*<br>(flatworm)] | 62 | 6e-14 | 29.41 | CDI96766 |
| 4 | 5789-2454 | 1111 | DNA polymerase | TPA_asm PolB<br>[Tasmanian devil<br>feces bidnavirus 1] | 55 | 9e-58 | 31.35 | DAC80328 |
| 5 | 6538-5789 | 249 | Uridine<br>diphosphate<br>glucose<br>pyrophosphatase | No blast hit | n/a | n/a | n/a | n/a |
|  | 6023-6571 | n/a | Inverted repeat | n/a | n/a | n/a | n/a | n/a |

### 3.4 Phylogenetic analysis revealed a new Bidnavirus in *Mr*

Because the Family *Bidnaviridae* includes viruses that appear to have evolved by the fusion of genetic elements from two different viral families (Krupovic et al., 2014; Krupovic and Koonin 2016), it is not possible to construct sensible phylogenetic trees using the whole genome sequence of MHBV as the basis for determining relationships. Only the single PolB domain of the MHBV genome gave any significant similarity to any known virus proteins in the current GenBank database. All its other putative open reading frames gave no significant hits. In addition, many hits with high similarity to the MHBV-PolB domain were listed as viruses in the Family *Parvoviridae*, a family that does not include viruses with DNA polymerase genes. Thus, it is apparent that current GenBank names for various viruses cannot be used to indicate their true phylogenetic position with respect to MHBV. Furthermore, a number of the GenBank records have arisen from DNA samples obtained from vertebrate animal feces (e.g., birds and mammals). This further complicates the problem because the originating viral DNA may have arisen from parasites or eaten prey rather than the source animal itself. To avoid these problems, we have focused solely on the region of the MHBV genome with homology to the PolB domain.

An alignment of deduced amino acid sequences from the top 8 hits for the PolB domain of MHBV is shown in **Fig. 4** and percent identity indices (31.6 to 34.6% for MHBV) are shown in **Table 3**. In addition, a cladistic tree based on the top 18 hits for the deduced protein sequence of the MHBV-PolB domain is shown in **Fig. 5**. Despite the fact that some of the GenBank records are given as parvoviruses, none of such hits could be from true parvoviruses.

**Table 3.** Percent identity matrix for the alignment in Fig. 5, created by Clustal2.1.

|  |  |  |  |  |  |  |  |  |  |  |
| --- | --- | --- | --- | --- | --- | --- | --- | --- | --- | --- |
| 1: | QQL94243_Macrobrachium | 100.00 | 32.61 | 31.56 | 34.20 | 34.12 | 33.26 | 34.19 | 34.75 | 34.55 |
| 2: | AKU37609_Bombyx_bidna | <u>32.61</u> | <u>100.00</u> | 38.45 | 41.63 | 38.82 | 38.95 | 42.19 | 40.00 | 40.21 |
| 3: | QTE03832_Lanius_denso | 31.56 | <u>38.45</u> | <u>100.00</u> | 39.57 | 42.41 | 41.29 | 42.20 | 39.83 | 40.25 |
| 4: | DAC80328_Tasmanian_bidna | 34.20 | 41.63 | <u>39.57</u> | <u>100.00</u> | 44.99 | 44.04 | 45.63 | 41.70 | 42.74 |
| 5: | QTE03818_Parus_denso | 34.12 | 38.82 | 42.41 | <u>44.99</u> | <u>100.00</u> | 82.06 | 72.32 | 71.77 | 73.02 |
| 6: | QVW56804_Phoenicurus_parvo | 33.26 | 38.95 | 41.29 | 44.04 | <u>82.06</u> | <u>100.00</u> | 72.58 | 72.84 | 73.08 |
| 7: | QTE03843_Passer_parvo | 34.19 | 42.19 | 42.20 | 45.63 | 72.32 | <u>72.58</u> | <u>100.00</u> | 73.99 | 75.66 |
| 8: | QTE03994_Motacilla_denso | 34.75 | 40.00 | 39.83 | 41.70 | 71.77 | 72.84 | <u>73.99</u> | <u>100.00</u> | 80.57 |
| 9: | QKK82936_Sonchus_parvo | 34.55 | 40.21 | 40.25 | 42.74 | 73.02 | 73.08 | 75.66 | <u>80.57</u> | <u>100.00</u> |

**Figure 4.**
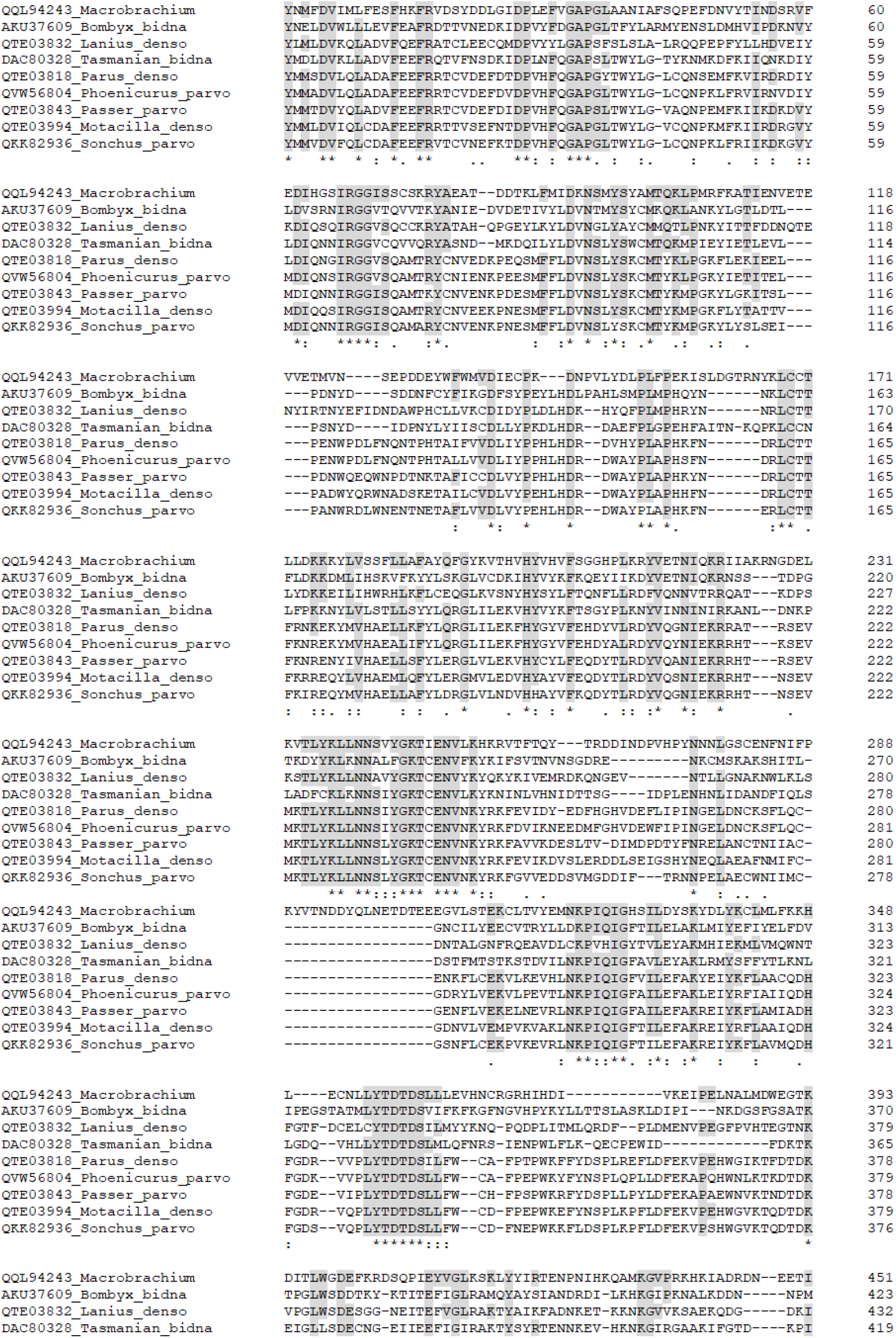
CLUSTAL Omega (1.2.4) multiple sequence alignment of 8 top blastp hits for the translated *PolB* domain in the MHBV genome.

**Figure 5.**
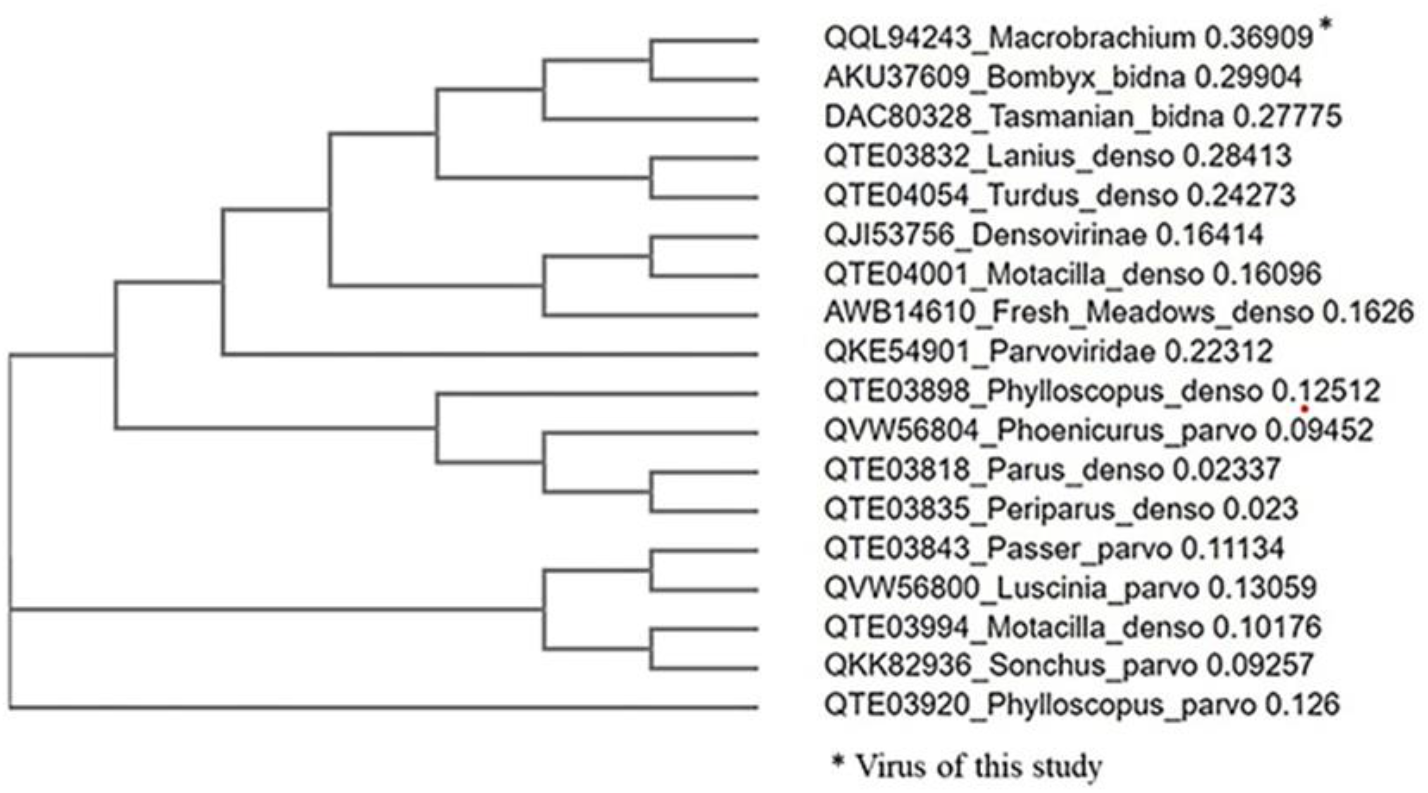
CLUSTAL Omega (1.2.4) cladistic analysis of 18 top blastp hits for the deduced protein sequence of the *PolB* domain in the MHBV genome. Aside from the three records at the top of the tree listed as bidnaviruses, all others were listed as parvoviruses or densoviruses, which must be incorrect, since the sequences are for PolB, which does not occur in parvoviruses.

### 3.5 ISH confirmed MHBV presence in normal nuclei and in intranuclear inclusions

The presence of MHBV was confirmed by ISH assays with tissue sections from our samples from 2009 and 2016 using a probe prepared with our amplified MHBV genomic DNA as the template. An unrelated probe using a fish virus DNA sequence was used as a negative control. This was better than using a no-probe control because it would reveal any non-specific DNA binding such as that which sometimes results, for example, with chitin in the crustacean cuticle. Samples from this study gave positive ISH signals (dark brown color) in the magenta intranuclear inclusions and in grossly normal nuclei in the HP (**Fig. 6**). This result revealed that the viruses in the 2009 and 2016 samples were most likely the same. In addition, this result indicated a possible explanation for positive ISH reactions in normal nuclei but lack of positive ISH signals in intranuclear inclusions in an earlier Thai report on detection of a type of DHPV in *Mr* samples (Srisala et al., 2021) (**Fig. 7**). This would be that the *Mr* PL in their study had dual infections of DHPV and MHBV. To test this hypothesis, archived DNA from the study of Srisala et al., 2021 was subjected to PCR testing methods for both DHPV and MHBV.

**Figure 6.**
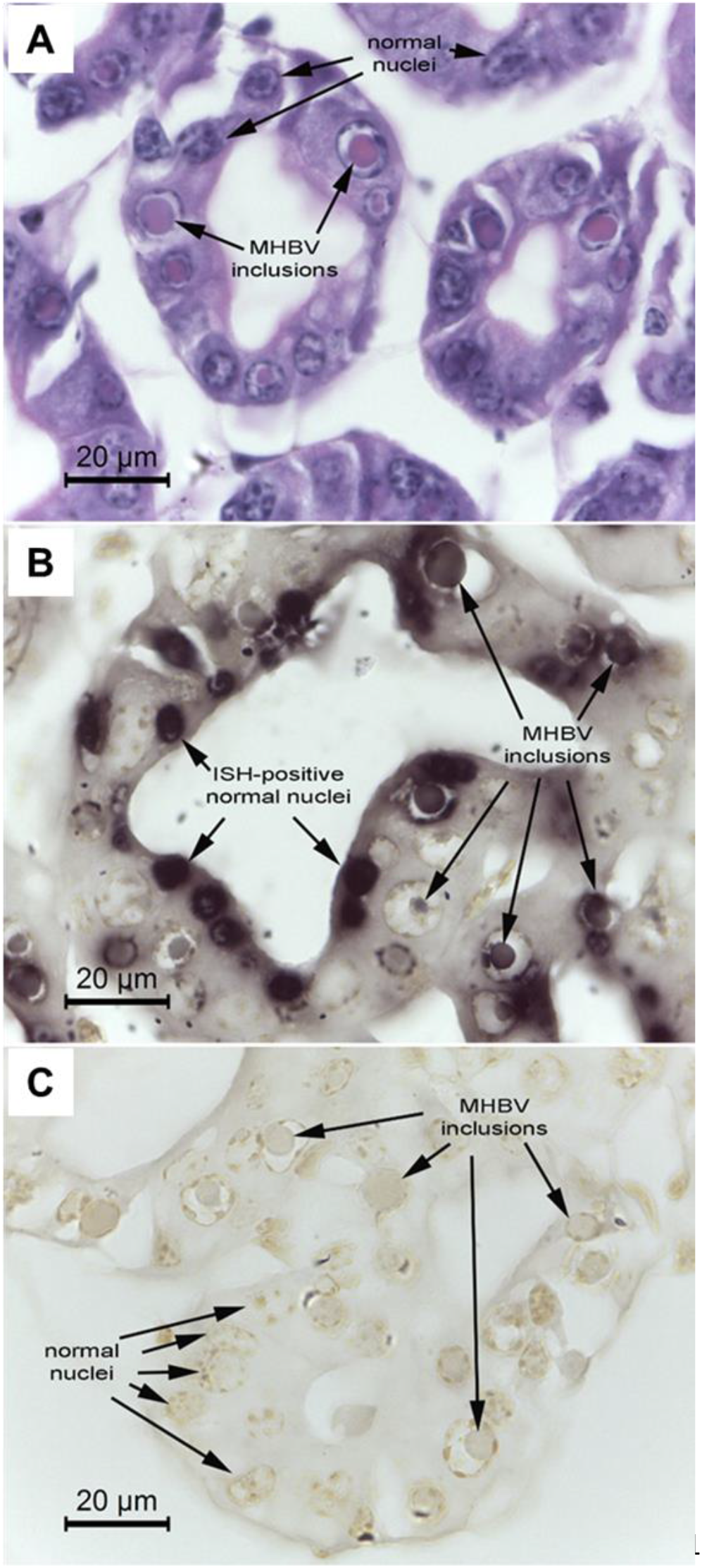
Example photomicrographs of ISH detection of MHBV in HP tissue using an MHBV-specific probe. (A) H&E-stained tissue section showing MHBV eosinophilic intranuclear inclusions. (B) Positive ISH reactions (dark brown staining) in both MHBV inclusions and in normal nuclei. Note that the inclusions give less intense ISH reactions than the infected, grossly normal nuclei. (C) Negative control probe (TiLV DNA) showing no positive ISH reactions.

**Figure 7.**
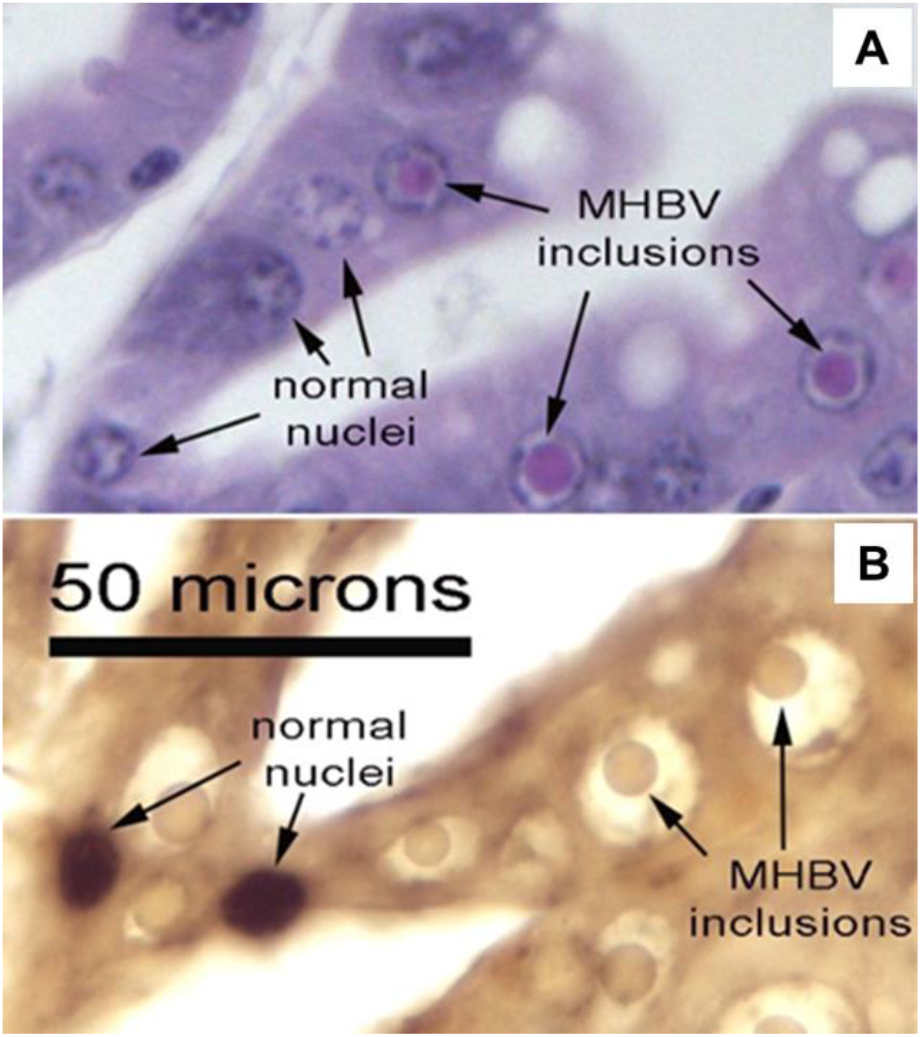
Example photomicrographs like those previously published (Srisala et al., 2021) and included here for comparison to Fig. 6. (A) H&E-stained tissue section showing MHBV eosinophilic intranuclear inclusions. (B) Previous result of Srisala et al. showed intranuclear inclusions gave positive ISH results for only DHPV (dark brown staining) with grossly normal nuclei and not with MHBV intranuclear inclusions.

### 3.6 Dual infections in the Srisala et al. study confirmed by PCR

Ten archived DNA extracts from the study by Srisala et al., (2021) (Sample Set 2), were subjected to PCR detection using the universal DHPV (DHPV-U) nested PCR method described in their study together with the MHBV one-step PCR method described herein. All ten samples gave positive amplicons for both methods, indicating dual infections of DHPV and MHBV (**Fig. 8**). The *Mr* actin gene was used as an internal control. At the same time, pooled DNA from 12 PL samples from our primary samples (Sample Set 3) gave positive PCR amplicons for MHBV only.

**Figure 8.**
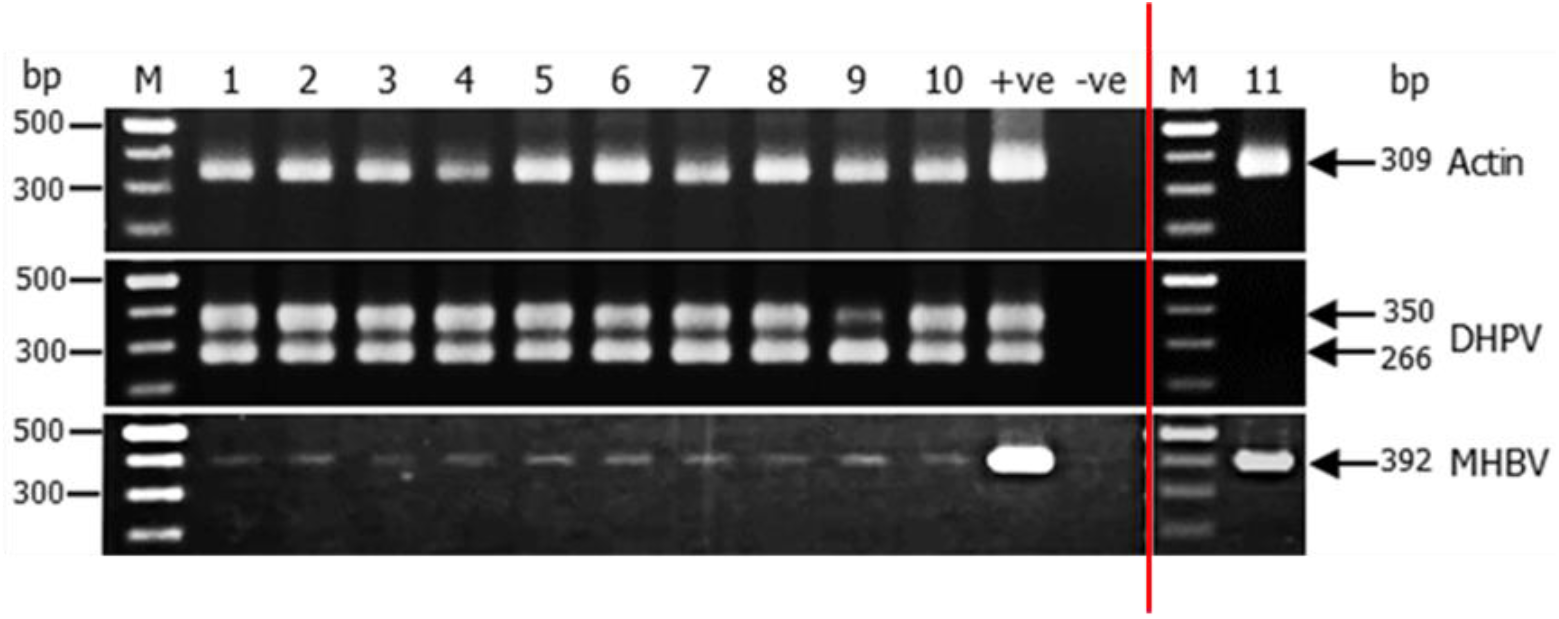
Agarose gels showing positive PCR test results for both DHPV and MHBV in archived DNA of 10 samples from the study of Srisala et al. (2021) (left) but positive PCR test results for MHBV only in 6 pooled DNA samples from the primary specimens in this study (right).

### 3.7 MHBV and DHPV in *Mr* are independent viruse*s*

Our PCR and ISH test results described in Section 3.3 and 3.4 revealed that the magenta intranuclear inclusions in *Mr* arose from MHBV, while the ISH results described in Section 3.5 revealed that the *Mr* specimens in the study of Srisala et al. (2021) had dual infections. To test this concept further using the DHPV and MHBV PCR methods, an additional 43 samples of grossly normal PL were taken in May 2022 (Sample Set 4) from the same hatchery as our primary Sample Set 3. Lightly positive results for DHPV were obtained with samples No.27 and 28 (2/43, 5%), and a lightly positive result was obtained for MHBV with sample No. 20 (1/43, 2%) (data not shown). None of the samples gave dual positive results. Since the dual infections of Srisala et al., 2021 were associated with mortality while the single infections with MHBV were not (including those of Sample Sets 3 and 4), it is possible that dual infections of DHPV and novel MHBV are required to cause *Mr* mortality. This possibility requires further investigation using bioassays with the respective purified viruses. If the occurrence of dual infections is a random occurrence, a prevalence of 0.05 for DHPV and 0.02 for MHBV would suggest that the probability of a dual infection would have been 0.001 at the time the samples were taken in 2022.

A similar situation previously existed regarding the relationship between the expression of white-tail disease (WTD) in *Mr* with respect to the dual infections with M. rosenbergii nodavirus (MrNV) and extra-small virus (XSV) that occurred together in WTD shrimp. XSV is a satellite virus (Qian et al., 2003; Sri-Widada and Bonami, 2004) that is dependent on the MrNV RNA-dependent RNA polymerase (RdRp) for its replication. Eventually, investigations revealed that some WTD shrimp were infected with MrNV only (Yoganandhan et al., 2006). In addition, separation, and testing of XSV and MrNV supported pathogenicity for MrNV alone (Zhang et al., 2006), as did pathogenicity of an MrNV infectious clone (Gangnonngiw 2020).

The source hatchery for the study by Srisala et al. (2021) (Sample Set 2) and for this study (Sample Sets 3 and 4) was the same, and that hatchery has been using the DHPV-U PCR method to clean its stock of DHPV. Now PCR testing for MHBV has been added to their screening procedures with the aim of eliminating both DHPV and MHBV from their domesticated *Mr* stock.

The MHBV genome sequence was submitted to GenBank under accession no. MN714903. The phylogenetic analysis and the presence of a PolB gene reveal that MHBV is not in the family *Parvoviridae* that includes the new sub-family *Hamaparvovirinae* (i.e., with the genera *Hepanhamaparvovirus* and *Penstylhamaparvovirus* that are common in penaeid shrimp species) (Pénzes et al., 2020). DHPV that sometimes occurs together with MHBV would clearly fall within the genus *Hepanhamaparvovirus*. The Family *Parvoviridae* is now divided into 3 sub-families, the *Parvovirinae* and *Densovirinae* and *Hamaparvovirinae*. Members of the sub-family *Parvovirinae* infect vertebrates (mammals, birds, and reptiles) and members of sub-families *Densovirinae* and *Hamaparvovirinae* infect invertebrates (insects, crustaceans, and echinoderms) (Pénzes et al., 2020; Cotmore et al., 2019). These proposals were accepted by the ICTV in March 2020 (Pénzes et al., 2020).

## 4. SUMMARY

We have discovered MHBV as the cause of eosinophilic to magenta, intranuclear inclusions in the hepatopancreas of *Mr* PL that are sometimes associated with mortality and sometimes not. DHPV, on the other hand showed no recognizable lesions in our samples, although grossly normal nuclei were shown to be infected with it by ISH. At the same time, PCR testing revealed that the two viruses can occur independently in PL samples that show no signs of disease. On the other hand, one set of samples exhibiting mortality was proven to have dual infections of MHBV and DHPV (Sample Set 2), raising the possibility that PL mortality occurs only with dual infections.

Based on the similarity of its ssDNA genome structure and a DNA polymerase B domain homologous to that of Bombyx mori bidensovirus V1, we believe that MHBV should currently be classified as a new member of the family *Bidnaviridae* with a genome consisting of a single ssDNA fragment.

Although MHBV sometimes occurs together with the true parvovirus DHPV, the current evidence suggests that they are independent ssDNA viruses. The fact that MHBV can exist with a single ssDNA strand, suggests to us that flacherie disease of silkworms may also be a dual infection with two viruses that belong to different families. This is supported by the fact that V1 and V2 of BmBV do not homologous each have complementary plus and minus ssDNA pair for each of has their own virion with a distinct capsid protein. To test this proposal for independent viruses, the virions containing the V1 and V2 fragments would have to be separated and used to challenge silkworm larvae independently and together. If they can replicate independently, they would be independent viruses whether they need to be together to cause flacherie disease or not. Similar bioassays are needed to determine whether dual infections of MHBV and DHPV are required to cause mortality in *Mr*.

A key feature for MHBV is the occurrence of the highly conserved DNA *PolB* found in ORF4 (position 5789-2454) along with an inverted repeat (position 7-126) (**Table 1)**. It has been suggested that an ancestral parvovirus genome was integrated into a large virus-derived DNA transposon of the Polinton/Maverick family (polintoviruses) to give rise to bidnaviruses (Krupovic et al., 2014; Krupovic and Koonin 2016). The alternative scenario proposing the origin of bidnaviruses from a parvoviral ancestor that acquired the *PolB* gene and TIRs from polintoviruses via horizontal gene transfer was considered less feasible (Krupovic and Koonin 2014b).

If the family name *Bidnaviridae* and the genus name *Bidensovirus* refer to an origin by horizontal fusion of a polintovirus ancestor and a parvovirus ancestor or some other virus family ancestor, then MHBV may fit the description of genus *Bidensovirus*. This would require data indicating that a major functional part of the MHBV genome arose from a parvovirus or other virus line. Our somewhat superficial analysis did not reveal such a link. However, if *Bidnaviridae* defines a family with a genome that consists of 2 distinct and inter-dependent ssDNA fragments, each with a different evolutionary past, then MHBV would not fit the description.

## CRediT authorship contribution statement

**Warachin Gangnonngiw**: Conceptualization, Methodology, Investigation, Data analysis, Writing original draft, and Funding acquisition. **Malinee Bunnontae**: Investigation and Data analysis. **Pattanapon Kayansamruaj**: Data Curation and Data analysis. **Saengchan Senapin**: Conceptualization, Methodology, and Writing original draft. **Jiraporn Srisala**: Investigation and Resources. **Timothy W. Flegel**: Conceptualization, Supervision, and Writing-reviewing and editing. **Kanokpan Wongprasert**: Conceptualization, Resources, Supervision, and Writing-reviewing and editing.

## Acknowledgments

This research project was supported by Mahidol University (Fundamental Fund: Basic Research Fund: the fiscal year 2022, Grant no. BRF1-054/2565). The authors would like to thank Panudda Meenium for her technical assistance.

## References

Anderson, I. G., Law, A. T., Shariff, M., & Nash, G. (1990). A parvo-like virus in the giant freshwater prawn, Macrobrachium rosenbergii. Journal of Invertebrate Pathology, 55(3), 447–449. https://doi.org/10.1016/0022-2011(90)90093-l

Bell, T. A., & Lightner, D. V. (1988). A handbook of normal penaeid shrimp histology. World Aquaculture Society.

Bolger, A. M., Lohse, M., & Usadel, B. (2014). Trimmomatic: a flexible trimmer for Illumina sequence data. Bioinformatics, 30(15), 2114–2120. https://doi.org/10.1093/bioinformatics/btu170

Cotmore, S. F., Agbandje-McKenna, M., Canuti, M., Chiorini, J. A., Eis-Hubinger, A.-M., Hughes, J., Mietzsch, M., Modha, S., Ogliastro, M., Pénzes, J. J., Pintel, D. J., Qiu, J., Soderlund-Venermo, M., Tattersall, P., & Tijssen, P. (2019). ICTV Virus Taxonomy Profile: Parvoviridae. Journal of General Virology, 100(3), 367–368. https://doi.org/10.1099/jgv.0.001212

Flegel, T. W. (2006). Detection of major penaeid shrimp viruses in Asia, a historical perspective with emphasis on Thailand. Aquaculture, 258(1-4), 1–33. https://doi.org/10.1016/j.aquaculture.2006.05.013

Gangnonngiw, W., Bunnontae, M., Phiwsaiya, K., Senapin, S., & Dhar, A. K. (2020). In experimental challenge with infectious clones of Macrobrachium rosenbergii nodavirus (MrNV) and extra small virus (XSV), MrNV alone can cause mortality in freshwater prawn (Macrobrachium rosenbergii). Virology, 540, 30–37. https://doi.org/10.1016/j.virol.2019.11.004

Gangnonngiw, W., Kiatpathomchai, W., Sriurairatana, S., Laisutisan, K., Chuchird, N., Limsuwan, C., & Flegel, T. (2009). Parvo-like virus in the hepatopancreas of freshwater prawns Macrobrachium rosenbergii cultivated in Thailand. Diseases of Aquatic Organisms, 85, 167–173. https://doi.org/10.3354/dao02075

Gotz, S., Garcia-Gomez, J. M., Terol, J., Williams, T. D., Nagaraj, S. H., Nueda, M. J., Robles, M., Talon, M., Dopazo, J., & Conesa, A. (2008). High-throughput functional annotation and data mining with the Blast2GO suite. Nucleic Acids Research, 36(10), 3420–3435. https://doi.org/10.1093/nar/gkn176

Hu, Z., Li, G., Li, G., Yao, Q., & Chen, K. (2013). Bombyx mori bidensovirus: The type species of the new genus Bidensovirus in the new family Bidnaviridae. Chinese Science Bulletin, 58(36), 4528–4532. https://doi.org/10.1007/s11434-013-5876-1

Krupovic, M., Bamford, D. H., & Koonin, E. V. (2014). Conservation of major and minor jelly-roll capsid proteins in Polinton (Maverick) transposons suggests that they are bona fide viruses. Biology Direct, 9(1), 6. https://doi.org/10.1186/1745-6150-9-6

Krupovic, M., & Koonin, E. V. (2014). Evolution of eukaryotic single-stranded DNA viruses of the Bidnaviridae family from genes of four other groups of widely different viruses. Scientific Reports, 4(1). https://doi.org/10.1038/srep05347

Krupovic, M., & Koonin, E. V. (2016). Self-synthesizing transposons: unexpected key players in the evolution of viruses and defense systems. Current Opinion in Microbiology, 31, 25–33. https://doi.org/10.1016/j.mib.2016.01.006

Li, D., Liu, C.-M., Luo, R., Sadakane, K., & Lam, T.-W. (2015). MEGAHIT: an ultra-fast single-node solution for large and complex metagenomics assembly via succinct de Bruijn graph. Bioinformatics, 31(10), 1674–1676. https://doi.org/10.1093/bioinformatics/btv033

Lightner, D. V., Redman, R.M., Poulos, B.T., Mari, J.L., Bonami, J.R., Shariff, M. (1994). Distinction of HPV-type viruses in Penaeus chinensis and Macrobrachium rosenbergii using a DNA probe. Asian Fisheries Science, 7(4), 267–72.

Lu, K.-Y., Huang, Y.-T., Lee, H.-H., & Sung, H.-H. (2006). Cloning the prophenoloxidase cDNA and monitoring the expression of proPO mRNA in prawns (Macrobrachium rosenbergii) stimulated in vivo by CpG oligodeoxynucleotides. Fish & Shellfish Immunology, 20(3), 274–284. https://doi.org/10.1016/j.fsi.2005.05.001

Mari, J., Lightner, D., Poulos, B., & Bonami, J. (1995). Partial cloning of the genome of an unusual shrimp parvovirus (HPV):use of gene probes in disease diagnosis. Diseases of Aquatic Organisms, 22, 129–134. https://doi.org/10.3354/dao022129

Menzel, P., Ng, K. L., & Krogh, A. (2016). Fast and sensitive taxonomic classification for metagenomics with Kaiju. Nature Communications, 7(1). https://doi.org/10.1038/ncomms11257

Pénzes, J. J., Söderlund-Venermo, M., Canuti, M., Eis-Hübinger, A. M., Hughes, J., Cotmore, S. F., & Harrach, B. (2020). Reorganizing the family Parvoviridae: a revised taxonomy independent of the canonical approach based on host association. Archives of Virology, 165(9), 2133–2146. https://doi.org/10.1007/s00705-020-04632-4

Phromjai, J., Boonsaeng, V., Withyachumnarnkul, B., & Flegel, T. (2002). Detection of hepatopancreatic parvovirus in Thai shrimp Penaeus monodon by in situ hybridization, dot blot hybridization and PCR amplification. Diseases of Aquatic Organisms, 51, 227–232. https://doi.org/10.3354/dao051227

Qian, D., Shi, Z., Zhang, S., Cao, Z., Liu, W., Li, L., Xie, Y., Cambournac, I., & Bonami, J-R. (2003). Extra small virus-like particles (XSV) and nodavirus associated with whitish muscle disease in the giant freshwater prawn, Macrobrachium rosenbergii. Journal of Fish Diseases, 26(9), 521–527. https://doi.org/10.1046/j.1365-2761.2003.00486.x

Rice, P., Longden, I., & Bleasby, A. (2000). EMBOSS: The European Molecular Biology Open Software Suite. Trends in Genetics, 16(6), 276–277. https://doi.org/10.1016/s0168-9525(00)02024-2

Seemann, T. (2014). Prokka: rapid prokaryotic genome annotation. Bioinformatics, 30(14), 2068–2069. https://doi.org/10.1093/bioinformatics/btu153

Srisala, J., Thaiue, D., Sanguanrut, P., Aldama-Cano, D. J., Flegel, T. W., & Sritunyalucksana, K. (2021). Potential universal PCR method to detect decapod hepanhamaparvovirus (DHPV) in crustaceans. Aquaculture, 541, 736782. https://doi.org/10.1016/j.aquaculture.2021.736782

Umesha, K. R., Dass, B. K. M., Manja Naik, B., Venugopal, M. N., Karunasagar, I., & Karunasagar, I. (2006). High prevalence of dual and triple viral infections in black tiger shrimp ponds in India. Aquaculture, 258(1-4), 91–96. https://doi.org/10.1016/j.aquaculture.2006.04.003

Widada, J. S., & Bonami, J.-R. (2004). Characteristics of the monocistronic genome of extra small virus, a virus-like particle associated with Macrobrachium rosenbergii nodavirus: possible candidate for a new species of satellite virus. Journal of General Virology, 85(3), 643–646. https://doi.org/10.1099/vir.0.79777-0

Yoganandhan, K., Leartvibhas, M., Sriwongpuk, S., & Limsuwan, C. (2006). White tail disease of the giant freshwater prawn Macrobrachium rosenbergii in Thailand. Diseases of Aquatic Organisms, 69, 255–258. https://doi.org/10.3354/dao069255

Zhang, H., Wang, J., Yuan, J., Li, L., Zhang, J., Bonami, J., & Shi, Z. (2006). Quantitative relationship of two viruses (MrNV and XSV) in white-tail disease of Macrobrachium rosenbergii. Diseases of Aquatic Organisms, 71, 11–17. https://doi.org/10.3354/dao071011

